# Performance of low-threshold, population replacement gene drives in cage populations of the yellow fever mosquito, *Aedes aegypti*

**DOI:** 10.1101/2024.08.06.606841

**Authors:** Zachary J. Speth, David G. Rehard, Patricia J. Norton, Alexander W.E. Franz

## Abstract

*Aedes aegypti* is the predominant vector for arboviruses including dengue, Zika, and chikungunya viruses, which infect over 100 million people annually. Mosquito population replacement strategies in which pathogen-susceptible mosquitoes in the field are replaced by laboratory-engineered pathogen-resistant strains are novel genetic control measures to inhibit the spread of malaria or arboviral diseases in endemic regions. To suppress arbovirus transmission following this approach, a mosquito strain needs to be transgenically modified to express an antiviral effector molecule which is linked to a gene drive (GD) system to both inhibit viral replication in the mosquito and drive the engineered resistance trait throughout a wild-type population. As a proof-of-concept, we tested the performance of two single locus CRISPR-Cas9 based GD for *Ae. aegypti* population replacement in small cage populations over 12 generations. Starting from a low release threshold of 1:9 GD bearing males, we observed two GD constructs in which Cas9 was expressed from different promoters increase in frequency in all discrete, non-overlapping cage populations. By generation 12, 56-79% of mosquitoes in six cage populations had at least one GD copy. The allele frequencies of the GD increased from <5% at release to >50% by G7 post-release for the *nanos*-driven Cas9 GD and by G10 in populations harboring the *zpg*-driven Cas9 GD. Insertion and deletion mutation (indel) frequency was measured for each discrete generation in pooled samples from the six populations harboring GD. We found that populations with Cas9 expression under control of the *nanos*-promoter accumulated gene drive blocking indels (GDBI) at more than twice the rate of populations harboring the *zpg*-promoter driven GD. Both GD produced *de novo* mutations at similar rates, with a difference in selection being the primary cause of greater indel accrual in the *nanos*-driven GD populations. Our results demonstrate that two single-locus, CRISPR-Cas9-based homing GD located at an intergenic locus exhibit continuous super-Mendelian inheritance in populations of *Ae. aegypti*. We further analyze the effects of fitness cost on the stability of low-threshold CRISPR/Cas9 based GD in populations of *Ae. aegypti*. This study demonstrates the feasibility of low-threshold, single-locus Cas9 gene drives for *Ae. aegypti* population replacement.

## Introduction

*Aedes aegypti*, also known as the yellow fever mosquito, is the predominant vector for arboviruses including dengue (DENV), Zika (ZIKV) and chikungunya viruses (CHIKV), which infect over 100 million people annually. *Ae. aegypti* is endemic globally throughout the tropical regions of the world and is established in the south-eastern and south-western United States (Khan et al., 2020). The *Ae. aegypti* population distribution is anticipated to expand northwards and southwards in the Western hemisphere within the coming decades as a result of global warming (Kraemer et al., 2019). *Ae. aegypti* was the principle vector responsible for a recent major ZIKV epidemic in Central and South America, (O’Reilly et al., 2018), and in 2024 it is the primary mosquito vector behind record dengue disease caseloads in the Western Hemisphere, predominantly in Amazonia (Leandro et al., 2024; Clarke et al., 2024). The efficacy of conventional vector control approaches, primarily insecticide application and mosquito habitat removal, is hindered by factors which include the presence of circulating insecticide resistance alleles, emerging insecticide resistant mutations, and the adaptations of *Ae. aegypti* to urban habitats (Garcia et al., 2018; Love et al., 2023; Wilke et al., 2020).

Population modification gene drives (GD) linked to anti-pathogen effectors is an approach to introduce pathogen resistance traits into mosquito populations, resulting in decreased vectorial capacity of the target pathogens. Currently, anti-malarial effectors coupled to GD in *Anopheles* mosquitoes are under development and have been shown in laboratory settings to effectively replace malaria susceptible populations with parasite-refractory mosquitoes (Gantz et al., 2015; Carballar-Lejarazu et al., 2023). Antiviral effector transgenes conferring resistance to DENV, ZIKV and CHIKV utilizing engineered long dsRNA inverted-repeat effectors (Franz et al., 2006; Franz et al., 2014; Williams et al., 2020), microRNA arrays (Liu et al., 2021), small RNA arrays (Buchman et al., 2019), single-chain antibodies (Buchman et al., 2020), and hammerhead ribozymes (Mishra et al., 2016; Mishra et al., 2023) have previously been demonstrated to suppress virus replication and systemic infection in *Ae aegypti* to varying degrees. Coupling antiviral effectors to GD is a necessary step to introduce antiviral effector transgenes into wild populations of mosquitoes such as *Ae. aegypti* thereby protecting the transgene from being lost between generations due to selection (Reid et al., 2021).

Gene drives are genetic elements which bias their own inheritance to super-Mendelian ratios (Burt and Trivers 2006). CRISPR-Cas9 based GD bias inheritance through their homing endonuclease activity, causing the GD bearing allele to be copied to replace the wild-type allele on the homologous chromosome via homology-directed DNA repair (HDR) of Cas9-induced DNA double-strand breaks (Gantz et al., 2015; Hammond et al., 2016; Windbichler et al., 2011). Homing is the predominant mechanism of CRISPR/Cas9-GD in *Anopheles* mosquitoes (Gantz et al., 2015; Hammond et al., 2016). While homing rates are often very high in *Anopheles* spp., in some cases resulting in greater than 95% transmission bias (Gantz et al., 2015; Carballar-Lejarazu et al., 2020), lower homing rates have been observed in *Ae. aegypti* for similarly designed GD (Li et al., 2020; Reid et al., 2022; Anderson et al., 2024; Verkuijl et al., 2022). In addition to decreased homing rates, *Ae. aegypti* CRISPR/Cas9 GD may also bias inheritance from non-homing related drive due to chromosome loss affecting unrepaired target chromosomes originating from non-GD carrying gametes (Verkuijl et al., 2022). Split GD systems for *Ae. aegypti* have previously been tested (Li et al., 2020; Verkuijl et al., 2022; Anderson et al., 2024). Split GD systems are confinable but require high release ratios of transgenic to wild-type insects, along with continuous releases over a sustained period, which may span several months of real time (Li et al., 2020; Anderson et al., 2024; Smidler et al., 2023). In contrast, low-threshold, single-component CRISPR/Cas9 GD can achieve high rates of population invasion from a single release of male mosquitoes (Gantz et al., 2015; Carballar-Lejarazu et al., 2020; Carballar-Lejarazu et al., 2023). The development and study of low-threshold GD in *Ae. aegypti* has thus far been limited to a single study (Reid et al., 2022).

Low-threshold GD allow for the placement of both the GD components and antiviral effectors into a single, stable genomic locus which is permissive for high levels of transgene expression (Franz et al., 2014; Reid et al., 2022). Studies of GD within laboratory populations of mosquitoes have previously yielded new insights into the drive dynamics, particularly with respect to the selection of GD resistant alleles (Hammond et al., 2017). Small population studies additionally provide an empirical reference for comparison with contemporary models of GD.

We tested the performance of two low-threshold, population replacement, single component CRISPR/Cas9 GD within six small cage populations of *Ae. aegypti*. The single component GD in this study consist of a polycistronic transgene comprising of Cas9 and sgRNA expression cassettes, and a fluorescent marker as cargo (**Fig. 1A**). The two GD express Cas9 from either the *Ae. aegypti nanos*-(AAEL012107) or *zero population growth* (*zpg*)-promoter sequences and 3’UTRs (*inexin-4*; AAEL006726) and are inserted in an intergenic locus on the 3^rd^ chromosome (designated “Carb109”, or C109), which is permissive for stable antiviral effector expression (Franz et al., 2014; Reid et al., 2022). For reasons of simplicity, we will name our two GD transgenes from here on *nanos*-GD and *zpg*-GD. The study concludes with an analysis of the stability of low-threshold population replacement GD as a function of genotype-specific fitness costs. A model for homing endonuclease GD targeting an intergenic locus, implemented in MGDrivE (Sanchez et al., 2019), is parameterized with different fitness cost scenarios and compared to observations from the cage trial study. The paper concludes with an analysis of effective engineered solutions for a population replacement GD, which would circumvent the introduction of gene drive blocking indels (GDBI).

**Figure 1.**
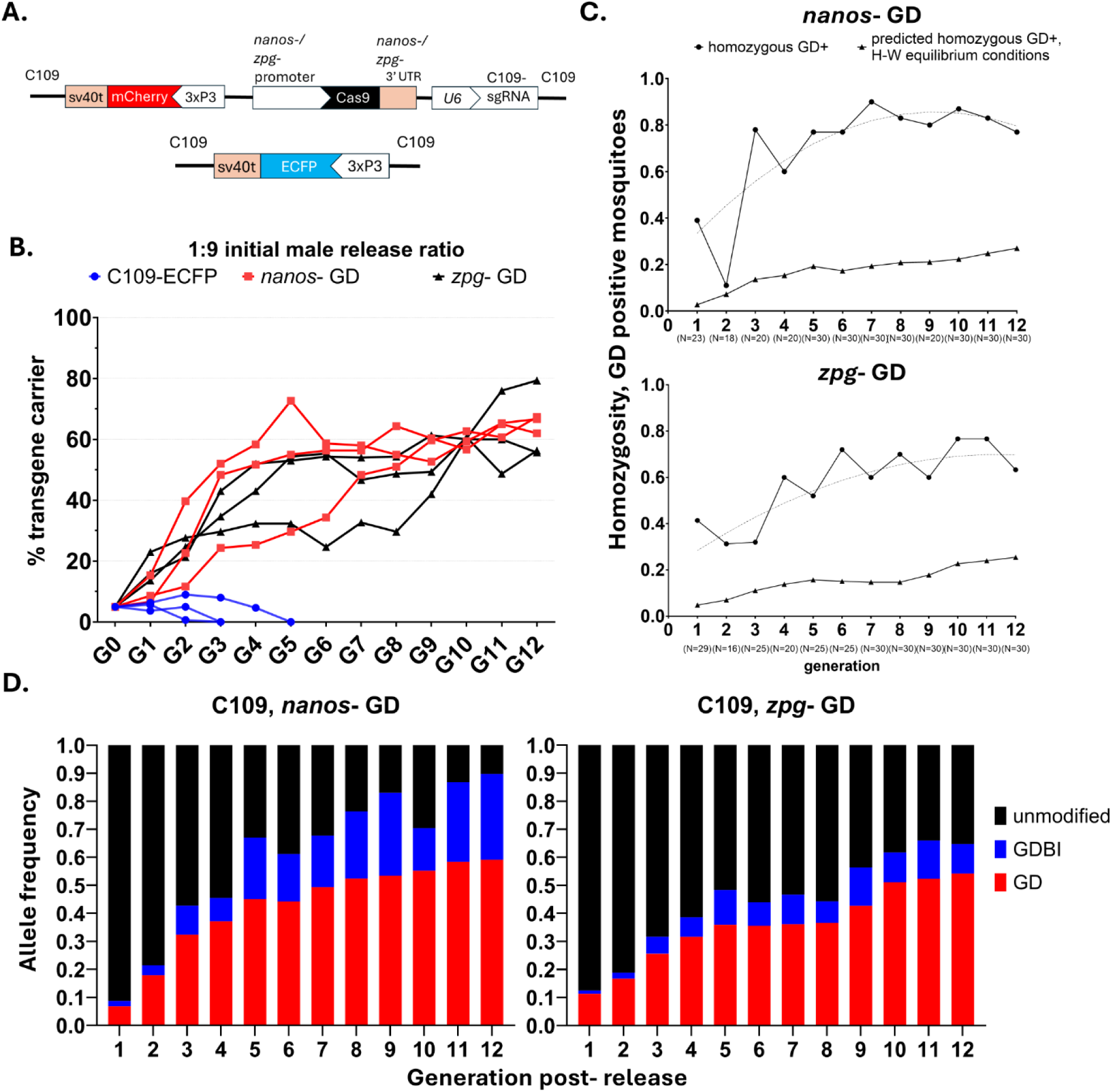
Single-locus CRISPR-Cas9 based GD exhibit continued super-Mendelian inheritance bias in small cage populations of *Aedes aegypti.* **A)** Schematic of the single-locus, CRISPR-Cas9 based GD transgene, and the control marker transgene at the *Carb109* locus (Reid *et al.,* 2022). **B)** Hemizygous GD carryingAe. *aegypti* males were released at a 1:9 ratio into populations of ’wild-type’ (HWE) *Ae. aegypti.* Non-overlapping populations with discrete generations were maintained in triplicate for 12 generations following the introduction of either *nanos-* or *zpg-* GD. Populations consisted of 300 individuals throughout cage trials. During pupation, 150 individuals of both sexes were randomly picked and analyzed for eye marker expression and placed into cages to start the next generation. **C)** GD carriers were genotyped using an allele-specific PCR test. Both *nanos-* and *zpg-* GD exhibited super-Mendelian inheritance and homing endonuclease activity, resulting in homozygote frequencies of GD carriers exceeding the expected frequency under Hardy-Weinberg equilibrium conditions in 11/12 generations for the *nanos-* GD and 12/12 generations for the *zpg-* GD. The *nanos-* GD exhibited greater rates of inheritance than the *zpg-* GD from generations 3 through 10 (binomial test, method of small p-values). **D)** Averaged allele frequencies from triplicate cage trial populations harboring GD alleles, unmodified alleles, and gene drive blocking indels (GOBI).

## Results

### Performance of two GD in small cage populations of *Ae. aegypti*

We chose a low cage release ratio of GD bearing hemizygous males to provide a sensitive test for GD performance over several generations. The GD designs are outlined in **Fig. 1A**. Starting with an initial 1:9 GD male to WT male release ratio (at G0), we observed a substantial increase in the number of GD carrying mosquitoes throughout three cage trial replicates for both the *nanos*-and *zpg*-GD drives at the C109 locus (**Fig. 1B**). *nanos*-GD carrying mosquitoes increased in proportion from 10% of the male carriers at release (G0) to >50% percent of the total population by G4 for two out of the three replicate populations. The *zpg*- GD exhibited a slightly delayed relative increase in GD carriers, reaching >50% invasion in two out of the three replicate populations by G5. By G12, however, 56-79% of the mosquitoes in all *nanos*-GD and *zpg*-GD populations had at least one GD copy. Strikingly, a similarly introduced control transgene containing a fluorescent marker cassette at C109 locus but lacking a GD was lost from all three replicate cage populations by G5. To test the hypothesis that C109 transgenic mosquitoes may have a baseline fitness deficit independent of the presence of the GD, we compared this result with Wright-Fisher models of genetic drift with no selection, starting with a conservative effective population size estimate. We found that the probability of transgene loss due to genetic drift was very low, even in populations with half the effective size of those used in our cage trials. This finding indicates a non-negligeable fitness cost associated with a transgene present at the intergenic C109 locus. We did not test whether the loss in fitness is a result of the fluorescent protein marker or is a result of inbreeding depression of the transgenic line, which was maintained independently in the laboratory for several generations.

### Homozygosity and allele frequencies

The allele frequencies of single-component GD at the C109 locus in the six cage populations was calculated from numerical recordings of GD carriers based on eye marker expression and their level of homozygosity based on allele specific PCR detection assays (**Fig. 1C**). GD positive mosquitoes in populations with both *nanos*- and *zpg*- GD exhibited ratios of homozygosity for the transgene far exceeding the expectations for a population meeting the conditions for Hardy-Weinberg equilibrium. The divergence in homozygosity for the GD alleles was maintained throughout the period of study (**Fig. 1C**). We observed a greater proportion of homozygous GD carriers in the *nanos*- GD than the *zpg*- GD populations from G3 onwards to G12 (**Fig. 1C**). The greater homozygosity of the *nanos*- GD may have resulted from increased homing rates, which is consistent with measurements for the *nanos*- GD in a single-cross study (Reid et al., 2022).

Between G3 and G10, 60-90% of *nanos*- GD (79.7% average) were homozygous. In comparison, from G3 onward to G10 samples from the *zpg*- GD individuals showed levels of homozygosity ranging from 32-77% (60.9% average), which was significantly less than the average nanos- GD carrier proportion (**Fig. 1C**). These observations indicate that the *nanos*-based GD construct exhibits a stronger overall drive. However, this data does not differentiate between a difference in GD performance due to either homing drive or meiotic drive mechanisms (Verkuijl et al., 2022).

Interestingly, the *nanos*- GD produced greater rates of homozygous offspring in relation to populations carrying the *zpg*-GD even as mating opportunities between GD carriers and naive individuals diminished while the predicted rates of random mating between GD carriers and individuals harboring GDBI increased.

### Sex-ratio of GD Inheritence

In the first generations after release, we observed for both GD a modest but significant sex bias for GD inheritance in males (**Figure 2A, B**). This sex bias waned over succeeding generations and was more persistent in populations with the *nanos*- GD, with the first three generations post release exhibiting a sex bias for male inheritance. There was no further evidence of sex-biased inheritance following G4 for the *nanos*- GD and following G2 for the *zpg*- GD (**Fig. 2B, Table S1**). This pattern may result from a bias for GD homing in male-forming zygotes or from a sex-dependent difference in viability associated with the GD transgene, which was lost after outcrosses to wild-type individuals within the small cage populations. It is important to note that the GD are not located on the same chromosome containing the sex-determining locus of *Ae. aegypti,* which is present on chromosome 1 (Hall et al., 2015; Aryan et al., 2020) (**Fig. 2C**).

**Figure 2.**
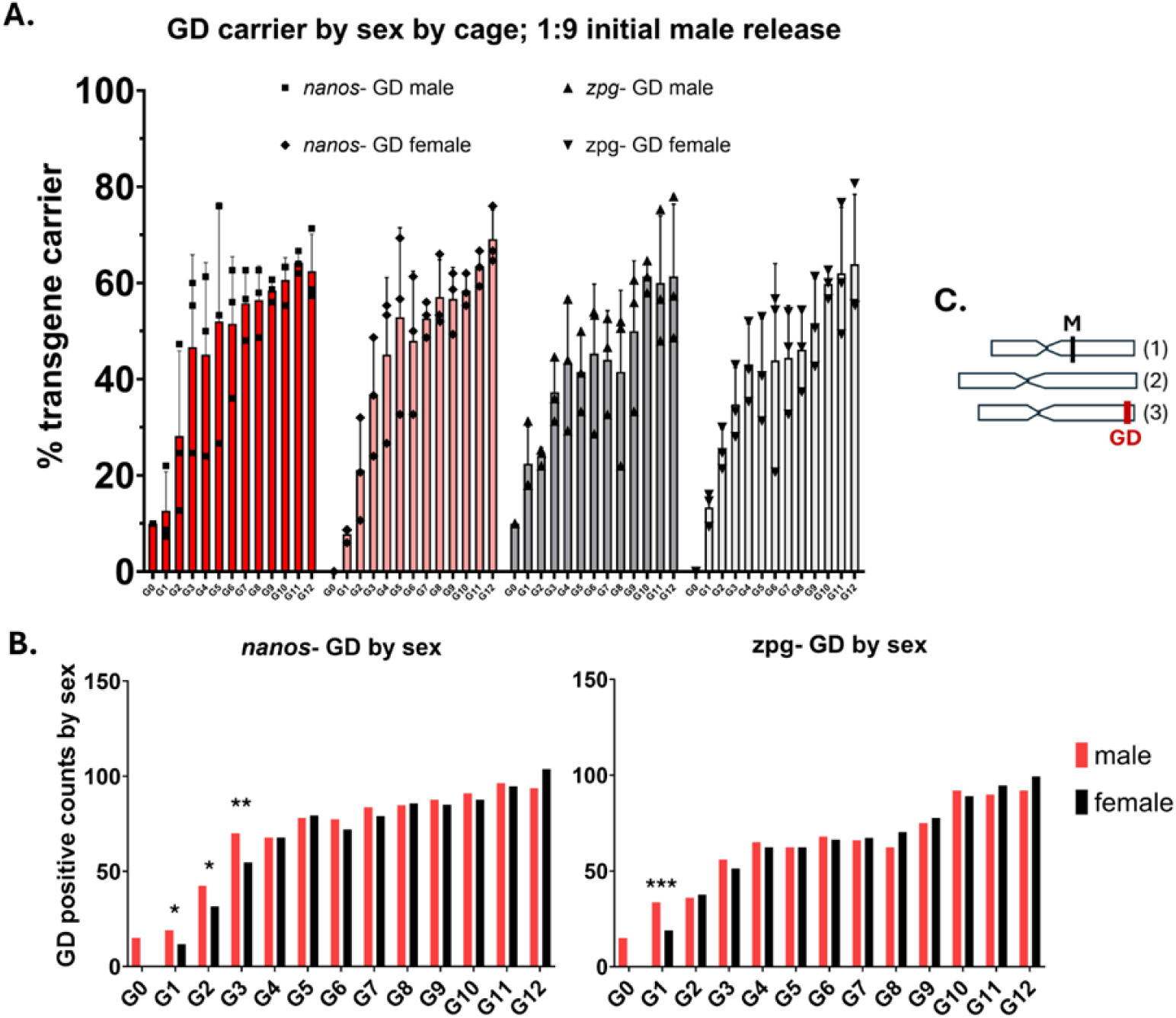
**A.** GD performances by sex in triplicate cage trial populations harboring *nanos-* and *zpg-* GD. B. Side by side comparisons of GD positive counts for male and female *Ae. aegypti* in populations with 300 mosquitoes each. A significant sex bias for increased male inheritance of the GD was detected from G1 through G3 in the nanos-GD populations and G1 in the zpg-GD populations (Fisher’s exact test, ’ p s 0.025, **ps .0025, *** p s .00025). Bars represent the averaged values of three replicates for counts of male and female GD carriers out of 150 individuals per cage. C. Schematic of Ae. *aegypti* chromosomes 1-3 with representative location of the male sex determining locus, Nix (marked M), on the first chromosome and the C109 GD target site on the third chromosome (marked GD).

### Assessment of Indels/GDBI

Deep sequencing results revealed striking differences in the accrual of gene drive blocking indels (GDBI) in populations with the *nanos*- or *zpg*- GD (**Figs. 1D, 3A, 4**). From the second generation onward *nanos*- GD carrying populations exhibited a greater proportion of GDBI at every generation than *zpg*- GD harboring populations. In addition, the proportions of GDBI as a share of non-GD alleles diverged over successive generations for the *nanos*- and *zpg*-populations (**Fig. 3A, Fig. 4**). By G12, the *nanos*- GD populations had accumulated GDBI in almost all of the target alleles which did not carry the GD, with *de novo* mutant alleles occupying >86% of non-GD alleles in two out of three *nanos*- GD populations (**Fig. 3A, Fig. 4**). In contrast, the accumulation of indels in *zpg*-populations was relatively muted, with mutant alleles occupying 23% on average of the non-GD alleles at G12 (**Figs. 2A, 3A**).

**Figure 3.**
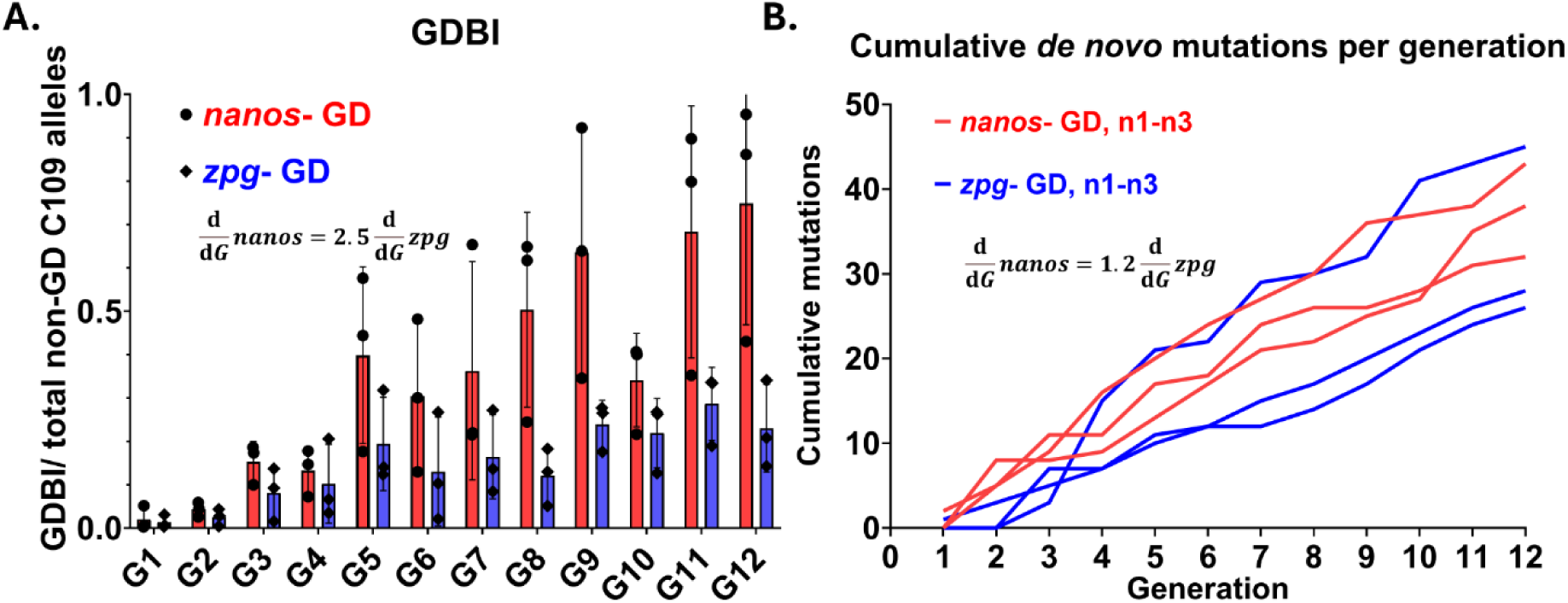
Promoter choice impacts insertion and deletion mutation accrualfor Aedes *aegypti* CRISPR/Cas9 GD. **A)** nanos-GD carrying populations exhibit a greater proportion of GD blocking indels (GDBI) than *zpg-* GD carrying populations. Total DNA was extracted from pooled samples of 100 larvae each, which were randomly collected for each cage trial replicate during generations 1-12 of the cage trial. A 537 bp amplicon spanning the GD target site was produced for 72 samples using PCR using with pooled genomic DNA. GDBI was measured by deep sequencing the pooled PCR amplicons spanning the GD target insertion site. B) Observed cumulative *de novo* mutations were counted for each cage trial population with *nanos-* or *zpg-* GD. Only mutations measured at frequencies greater than 0.5% of the reads per sample were considered in the analysis.

**Figure 4.**
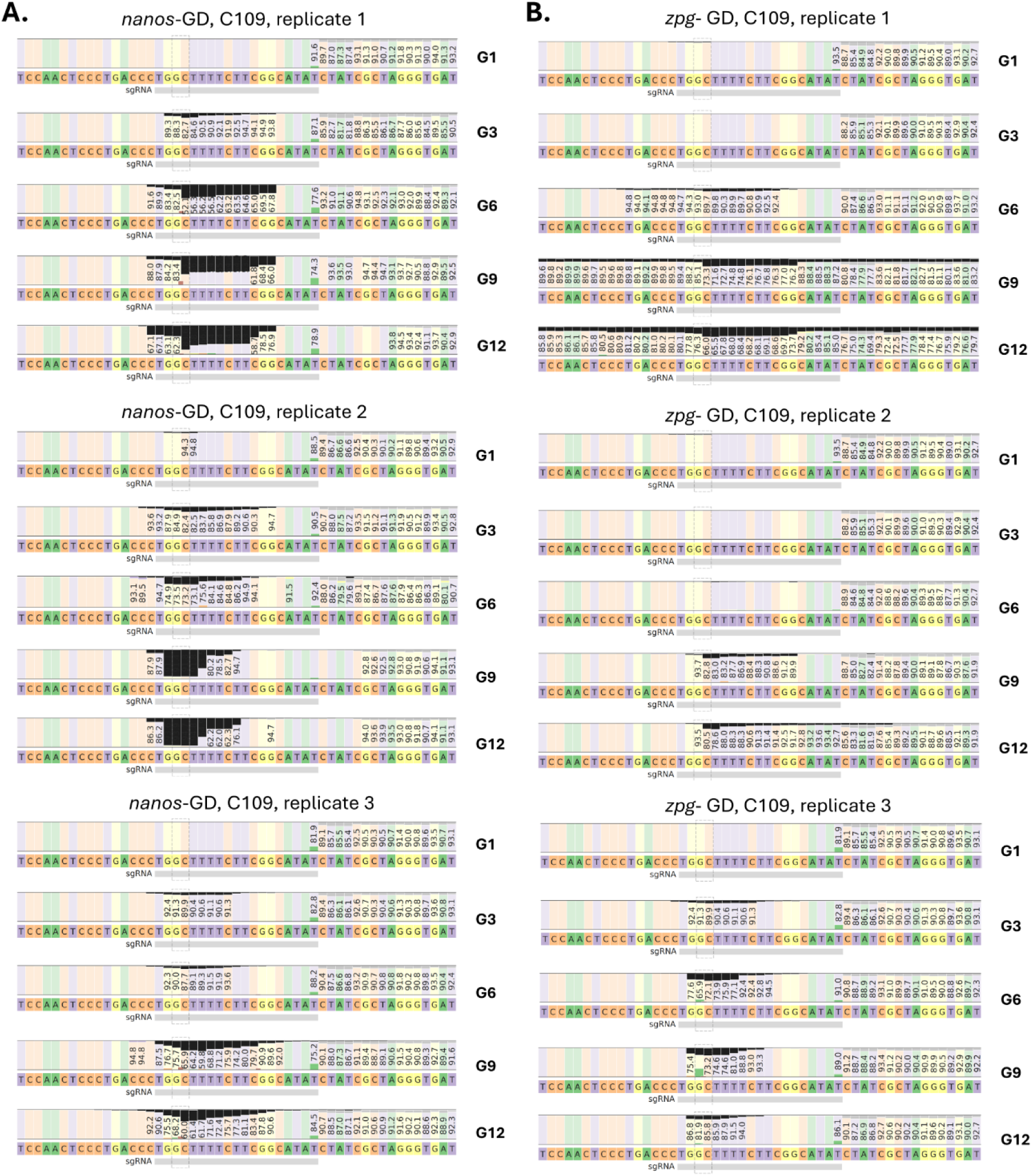
Evolution of GD blocking mutations at the sgRNA target site in representative generations of *Ae. aegypti small* cage populations harboring **A)** *nanos*- and **B)** *zpg*- GD. The fractions of aligned paired-end reads showing deletions at respective nucleotide positions are shown by black bars.

GDBI accrual could arise either due to the accrual of *de novo* mutations at the CRISPR-Cas9 target site as a result of double-strand DNA break repair employing a NHEJ mechanism, or from selection for GD resistance mutations. Distributions of indels from individual cage trial populations with the *nanos*- GD indicated selection and accrual of indels over time (**Fig. 4**). Although unique indels located at the intergenic C109 locus are unlikely to have significant differences in fitness cost, this pattern could arise from positive selection for GD resistance mutations arising within earlier generations in the population. Positive selection for GDBI at an intergenic target locus in *Ae*.

*aegypti* populations carrying a CRISPR/Cas9 homing endonuclease GD may occur due to shredding of chromosomes with unmodified alleles at the target site (Verkuijl et al., 2022). In order to determine whether the difference between GDBI accrual was more attributable to a large difference in rates of *de novo* mutations or was a result of positive selection, we counted the observed *de novo* indels for each population from 72 sets of paired-end reads of representative pooled samples from the GD carrying populations (**Fig. 3B**). In total, 120 unique *de novo* indels at the C109 locus were identified in reads from the six cage populations (**Table S2**). Mutations were predominantly short deletions of <15 base pairs pseudo-randomly distributed 5’ of the PAM, which is characteristic of NHEJ repair of Cas9-induced double-strand DNA breaks (Chauhan et al., 2023) (**Fig. 4, Table S2**). Several of the mutations were observed within multiple independent populations, including five short deletions at the target locus which were observed in all six of the cage trial populations (**Fig. S1**, **Table S2**). We found that for the *nanos*- GD populations, GDBI accrual exceeded the rates observed in *zpg*-populations by more than double (**Fig. 3A**). However, when comparing *de novo* mutation accumulation rates between populations with either GD construct, the difference was less than half of that observed for GDBI accrual as a fraction of non-GD alleles (**Fig. 3A, B**). These results indicated that the *nanos*- GD has 1) a significantly greater fitness cost, which is consistent with previous results from single-crosses which measured fertility and pupation rates (Reid et al., 2022), and 2) that differential selection for GD resistant alleles caused the substantial difference in observed accrual of GDBI over successive generations.

### Modeled behaviors of *nanos*- and *zpg*- GD

To further test the hypothesis that GDBI accrual in the *nanos*- GD populations results from an increased fitness cost to individuals with at least one GD allele and positive selection for GDBI, we modeled the performances of both *nanos*- and *zpg*- GD with different fitness cost parameters in MGDrivE (**Fig. 5**). The model utilizes a modified version of the GD function “Cube-CRISPR2MF.R”, which condenses all GDBI to a single allele type (Sanchez et al., 2019; Reid et al., 2022). In this scenario unique GDBI share the same fitness cost, which is expected to be the case for short deletion mutations at an intergenic locus. This GD model has been specified to our GD design principle and was parameterized with empirically determined measures of sex-specific GD homing rates, resistance allele formation rates, as well as maternal deposition rates for either the *nanos*- or *zpg*- GD (Reid et al., 2022).

**Figure 5.**
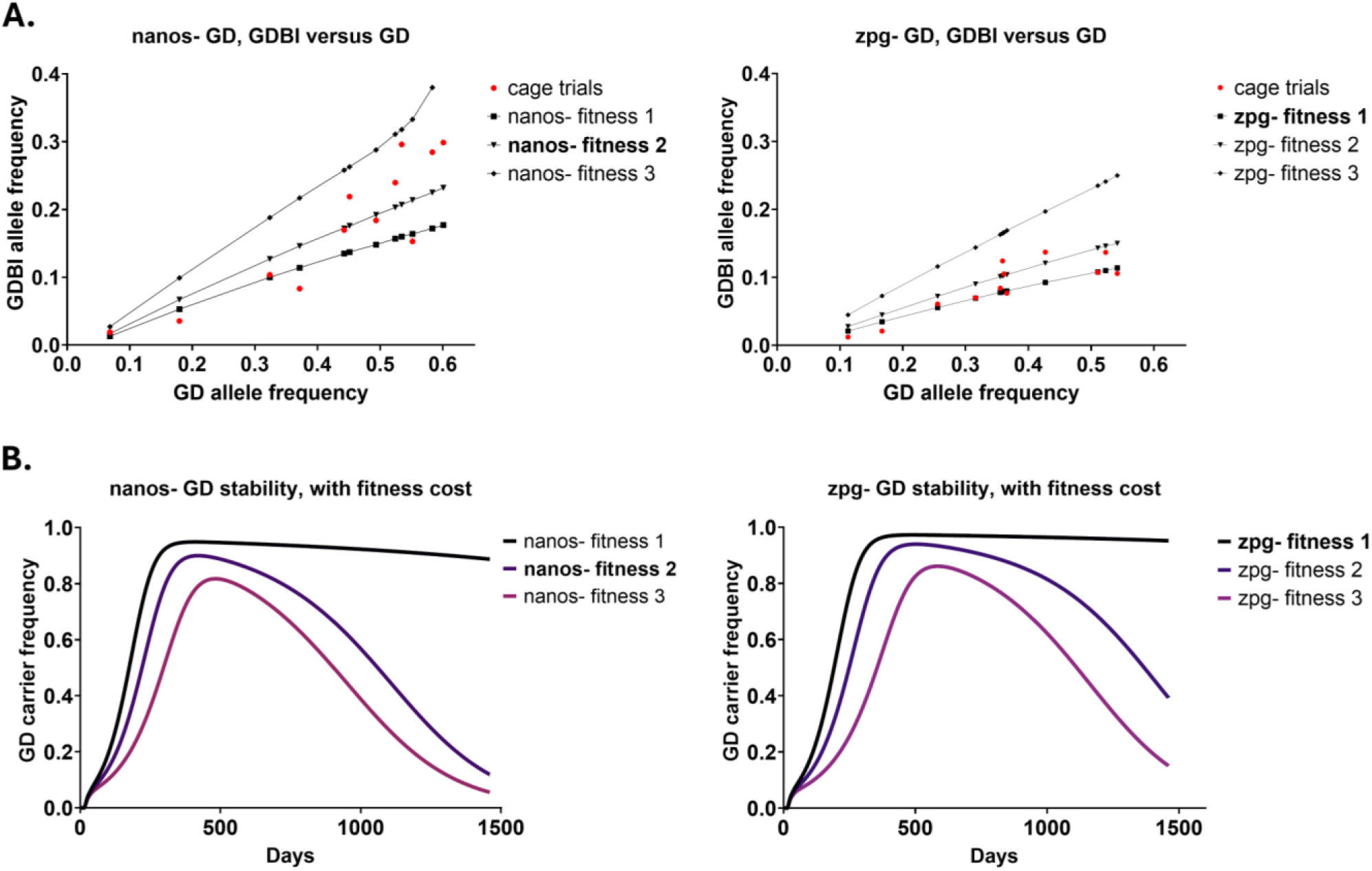
Observed versus modelled behavior of single-locus, population replacement GD with different fitness cost parameters. The model describes a single-locus CRIISPR/Cas9 based GD with sex-specific parameters for homing and resistance allele (GDBI) formation rates, and maternal deposition resulting in GDBI (Reid et al., 2022). The model is descriptive of a GD targeting an intergenic locus, in which the GD transgenes have an associated fitness cost, while GDBI alleles by default do not incur a fitness cost. The GD model with fitness parameters which most closely matches observations of GD behavior in cage trials is shown in bold. **A)** *nanos-* GD behavior most closely matched the model with fitness cost scenario 2, in which all individuals harboring a GD copy are penalized with a 10% reduction in fertility and a 10% reduction in pupation rate, *zpg-* GD behavior closely matches fitness cost scenario 1, in which GD carrying individuals have a 10% reduction in fertility without reduced pupation rates. B) Long term stability of GD under different fitness cost scenarios. Overall GD stability is highly sensitive to fitness costs of the carriers. The GD homing rates, resistance allele formation rates, and maternal deposition have a more modest but significant impact on long term GD-performance, as shown by comparisons between the *nanos-* and *zpg-* GD models with empirically determined parameters from previous single-cross studies (Reid et al., 2022),

The *nanos*- GD behavior in cage trials most closely matched the model with a 10% penalty to GD carrying individuals in both fertility and pupation rates, as determined by a minimal value of mean absolute scaled error in comparison with the model parameterized with lesser and greater fitness reductions of GD positive individuals (**Fig. 5A**, **Table 1**). The *zpg*- GD behavior most closely matched the GD model with a 10% reduction in fertility and no reduction in larval or pupal survival (**Fig. 5A**, **Table 1**). The fitness cost parameters for the models which best fit the cage trial results are in close agreement with measurements from single-crosses of *nanos*- and *zpg*- GD parents. In single-crosses with *zpg*- GD parents larval and pupal viability of GD carrying offspring was determined not to be significantly different from those values measured for the non-transgenic HWE recipient strain, while single-crosses with *nanos*- GD parents produced offspring with reduced larval viability (Reid et al., 2022).

**Table 1.** Fitness parameters and GD fitting to models. Fitness cost parameters for the various GD fitness scenarios shown in Figure 4.

| Model | genotype specific fitness parameters | days above 70% GD-carrier proportion |  | mean absolute scaled error of model versus observed values |  |
| --- | --- | --- | --- | --- | --- |
|  |  | <u>nanos- GD</u> | <u>zpg- GD</u> | <u>nanos- GD</u> | <u>zpg- GD</u> |
| fitness 1 | fertility "WH"="HH"="HR"=0.9 | 2297 | 3381 | 13.36 | **1.95** |
| fitness 2 | fertility and pupation rate "WH"="HH"="HR"=0.9 | 592 | 866 | **5.02** | 3.99 |
| fitness 3 | fertility "WH"="HH"="HR"=0.9, pupation rate "WH"=0.8,"HH"="HR"=0.9 | 339 | 485 | 11.07 | 16.86 |

Stability of population replacement GD over longer time periods is highly dependent on the fitness cost associated with the GD. The model is specific for a CRISPR/Cas9 based GD targeting an intergenic locus, the only difference in modeling scenarios tested was a change in the fitness cost parameters for genotypes carrying a GD. The models with the best ability to accurately forecast the GDBI alleles as a function of GD allele over intervals corresponding to a single cage trial generation have the lowest values of mean absolute scaled error and are bolded.

The model for our GD designs forecasts increased stability and invasion rates over longer time periods for the *zpg*- GD, which is supported by regression to observed allele frequencies in the cage trials (**Fig. 5A, B**). Although the *zpg*- GD exhibits lower homing rates, it outperforms the *nanos*-GD as a population replacement vehicle over longer time periods due to a combination of decreased mutation rates from NHEJ events and decreased fitness costs. Analysis of cage trial results and their comparison with the predictive power of GD-models with different fitness parameterizations indicates that the *zpg*- GD is expected to be stable as a vehicle for population replacement for between 2 ½ and 9 years, as defined by >70% invasion rate (**Table 1**). By this criterion, the *nanos*- GD is predicted to remain stable for a period 1 to 2 years before carrier frequency declines below the threshold (**Table 1**). Reduced fitness cost with selection for GDBI is the primary contributor to the increased stability over longer time intervals of the *zpg*- GD.

Decreased mutation rates from NHEJ repair of Cas9 induced double-strand DNA breaks of the *zpg*- GD in comparison to the *nanos*- GD more modestly contributes to the GD performance and stability (**Fig. 5B**).

## Discussion

Previous studies of CRISPR/Cas9 GD in *Ae. aegypti* have primarily focused on split GD systems to test the effects of promoter choice on GD homing rates, resistance allele formation and fitness cost (Li et al., 2020; Verkuijl et al., 2022; Anderson et al., 2023). GD function has been incrementally improved through the selection of germline specific promoters and multiplexing of sgRNAs (Anderson et al., 2024). Single-locus CRISPR/Cas9 based GD which achieve high rates of homing and drive the expression of linked antipathogen effectors have been described for population modification of *Anopheles* mosquitoes (Gantz et al., 2015; Carballar-Lejarazu et al., 2020; Carballar-Lejarazu et al., 2023). For *Ae. aegypti*, similar GD designs have been less vigorously pursued so far, since relatively low homing rates and relatively high rates of GDBI formation had been observed in earlier studies. However, while *Ae. aegypti* population *suppression* is unattainable with current CRISPR/Cas9 based GD performance characteristics (Marshall et al., 2017), robust and lasting population *modification* looks achievable in this mosquito species as our previous study (Reid et al., 2022) and the work here suggest. To our knowledge, this paper is the first study of low-threshold CRISPR/Cas9 based GD being evaluated in cage populations of *Ae. aegypti*. We describe two GD which can be coupled to deliver antiviral effectors in *Ae. aegypti* and which are capable of invading a large proportion of small, fixed mosquito populations, starting from a low threshold release. While neither GD will reach complete fixation due to indel accrual, both GD reached penetrance of >60% from an initial low release ratio. Our models of *Ae. aegypti* population replacement GD show that even when generating resistance at appreciable rates the GD may remain stable at relatively high invasion rates for multiple years (**Figure 5**). Extrapolating the *zpg*- GD performance observed in cage trials to GD models with different fitness costs indicates that the construct may remain stable at >70% population invasion for a period of 2.5-9 years, before being lost due to selection. Future efforts to develop GD for population modification strategies in *Ae. aegypti* should therefore aim to achieve greater GD stability primarily through fitness cost reduction.

GD function in host organisms varies according to differences in the utilization of DNA repair mechanisms during gametogenesis. In *Ae. aegypti*, a primary obstacle to GD development has been the formation of GDBI, which result from indel formation during repair of the Cas9-catalyzed double-strand (ds)DNA break by NHEJ or by single-strand annealing repair mechanisms (Basu et al., 2015; Bhargava et al., 2016; Finney et al., 2022). NHEJ predominates over HDR outside of a constrained temporal period during meiosis. This results in very low rates of donor directed repair during gene editing experiments in *Ae. aegypti* embryos, absent any targeted inhibition of NHEJ (Basu et al., 2015). Given the importance of the dsDNA break repair mechanism for accurate homing GD function, previous studies of CRISPR/Cas9 based GD in both *Anopheles* spp. and

*Aedes* spp. have investigated the effects of promoter choice on GD homing efficiencies and resistance allele formation (Li et al., 2020; Verkuijl et al., 2022; Hammond et al., 2021; Reid et al., 2022; Anderson et al., 2023). For *Ae. aegypti*, it should be noted that except for two of the studies (Li et al., 2020; Reid et al., 2022), the GD transgenes containing different promoters were integrated into different chromosomal loci due to transposon mediated transformation resulting in a quasi-random choice of the transgene integration site (Verkuijl et al., 2022; Anderson et al., 2023). This is important as position-effects in *Ae. aegypti* are known to contribute to transgene instability (Jasinskiene et al., 1998; Franz et al., 2009; Franz et al., 2014), (Reid et al., 2022), and may result in changes in GD expression/function (Verkuijl et al., 2020). The GD used in this study had been previously constructed using *Ae. aegypti nanos*-(AAEL012107) or *zpg* (*inexin-4*; AAEL006726) transcriptional regulatory sequences for Cas9 expression (Reid et al., 2022). The target locus for both GD, designated C109 (“Carb109”) (Franz et al., 2014), is located downstream of the distal-most protein encoding sequence (AAEL010318) on chromosome 3 (Reid et al., 2022). The C109 locus has previously been shown to be a genetically stable integration site for antiviral effectors (Franz et al., 2014). Our Cas9 promoter choice was based on previously revealed gene expression patterns and the GD design for *Anopheles* mosquitoes (Carballar-Lejarazu et al., 2020; Hammond et al., 2021). *Nanos*-transcripts are expressed in the female germ tissue of *Ae. aegypti*; they are produced by nurse cells within the mosquito oocyte, and are present within the germ cells during the earliest stages of embryonic development (Calvo et al., 2005; Adelman et al., 2007; Terradas et al., 2022). In contrast, *zpg*-transcripts are not maternally deposited, but are expressed within the germline cells primarily during the earliest stages of embryonic development (Hammond et al., 2021 Plos Genetics).

Standing allelic diversity at the sgRNA target site is another important consideration for GD design. Standing genetic variation at the target locus mitigates population suppression GD as a result of selection (Schmidt et al., 2020). Population replacement GD with low fitness cost are tolerant of standing resistance alleles, but exhibit a reduced level of maximal invasion. The C109 target locus is conserved in the Liverpool and HWE strains of *Ae. aegypti*, which have origins from collections in West Africa in 1930s and Puerto Rico in the 1990s, respectively. We did observe a single GDBI allele at C109 in about 8-9% of target amplicon reads from the Orlando strain of *Ae. aegypti* (**Figure S5**). Altogether this data suggests that the C109 locus is highly conserved, although there may be low frequency standing GDBI at the homing site in some populations of *Ae. aegypti*.

GD fitness cost may arise from off-target activity of the GD, although results from a whole-genome sequencing study indicate that this is likely a rare scenario for CRISPR-Cas9 based GD in mosquitoes (Garrood et al. 2021). More likely, the fitness reduction is attributable to a greater loss of non-GD bearing gametes assorting with the GD. This may arise from shredding of the GD target allele, which may occur in the embryo of GD bearing heterozygotes, or throughout development in somatic tissues. This source of fitness reduction would result in an observed fertility decrease in outcrosses of GD to WT individuals, and is consistent with previous observations of a fertility reduction in the *nanos*-GD (AeaNosC109^GD^) but not so in the *zpg*-GD (AeaZpgC109^GD^) line (Reid et al., 2023). A GD inheritance bias resulting from shredding of non-GD target loci has been identified among CRISPR/Cas9 based GD carrying *Ae. aegypti* (Verkuijl et al., 2022). This mechanism incurs a genetic disadvantage at the heterozygous level, which would lead to increased observations of homozygosity of GD bearing individuals at the target allele, in addition to an increased homing endonuclease activity. Indeed, this may have contributed to the elevated ratios of homozygotes observed in later generations of our *nanos*- GD harboring cage populations.

Otherwise, the level of homozygosity exclusively achieved via GD homing would be expected to decline between generations due to random assortment of alleles during sexual reproduction. In addition, progressively fewer wild-type alleles as GD targets are becoming available in the population from one generation to the next (**Figure 1**).

As the GD fitness approaches the resistance allele fitness, the stability of the GD and its invasiveness both increase (**Figure 5**). The GD and GDBI alleles reach a stable equilibrium if there is no difference in fitness between the GD-bearing and wild-type alleles (**Figure S4**). If the GD allele fitness eclipses that of GDBI, the GD would become fixed in the population (**Figure 6, Table S3**).

**Figure 6.**
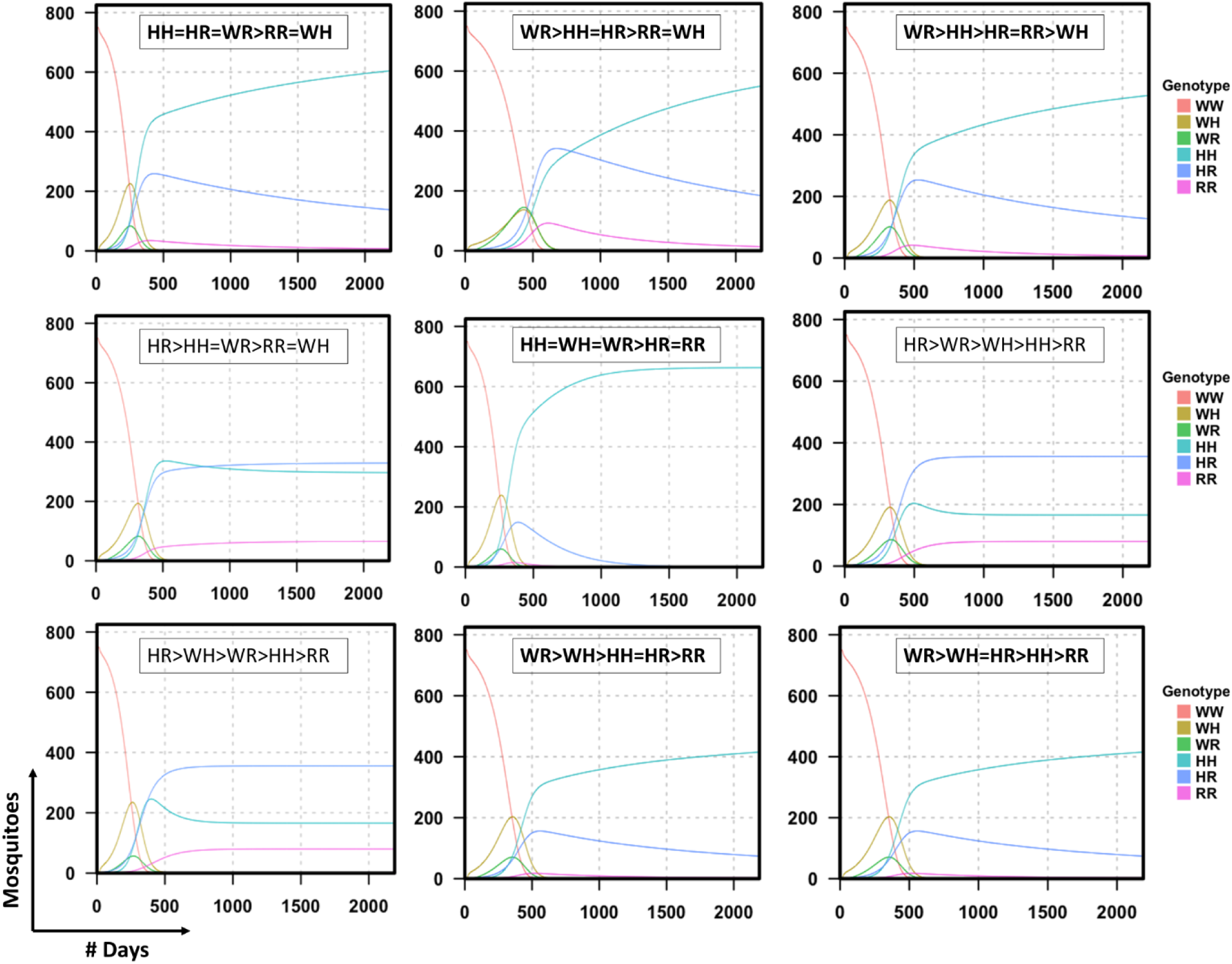
Equilibrium state and stability of low threshold population replacement GD at intergenic loci is determined by genotype-specific fitness inequalities. Deterministic simulations of low threshold population replacement GD were performed in MGDrivE using GD parameters which were empirically determined for the *nanos-* GD. The GD model is defined for a population replacement, single-locus CRISPR/Cas9 based GD with sex-specific homing and resistance allele formation and with maternal deposition (Reid et al., 2022). Different fitness inequalities with parameters listed in Table S3 were tested to determine design principles for population replacement GD, which reach fixation or stabLe equilibrium. In addition to the models shown in the figure, all GD with fitness inequalities HH AND WH > RR AND WR reach fixation. In **bold:** scenarios where HH reaches/ approaches fixation; not in bold: scenarios where a stable equilibrium state is reached between HH, HR and RR genotypes.

The fitness of alleles may vary in a genotype-specific manner. Different scenarios of allele fitness inequalities should therefore be considered when predicting GD dynamics in populations. The stability and equilibrium conditions of the GD and its resistance alleles are governed by the fitness inequalities between the genotypes in GD harboring populations (**Figure 6, Table S3**). Our GD models indicate that single-locus CRISPR/Cas9 GD for population replacement with the non-negligible GDBI formation rates observed for *Ae. aegypti* GD will invariably reach fixation or stable equilibrium if GD homozygotes (HH) are more fit than GD heterozygotes containing a resistance allele (HR) and RR homozygotes (**Figure 6, Table S3**). A more robust population replacement GD could thus be engineered by utilizing Cleave and Rescue (ClvR), or Home and Rescue (HomeR) GD designs (Oberhofer et al., 2019; Adolfi et al., 2020; Kandul et al., 2021).

HomeR and ClvR are designed to decrease the fitness of resistance alleles and simultaneously increase the fitness of the GD by targeting/disrupting and eventually restoring essential gene function. HomeR or ClvR targeting haplosufficient genes produce HH>RR fitness inequalities, thus biasing selection in favor of the GD. HomeR or ClvR GD targeting a haplo*in*sufficient gene are designed to produce WH>WR fitness inequalities in addition to WH>RR and HH>RR fitness inequalities but may be sensitive to the expression of the rescue transgene, which would reduce the fitness of HH genotypes in relation to WH genotypes thereby leading to decreased homing rates (Hou et al., 2024). Both ClvR and HomeR may be susceptible to functionally restorative mutations of the targeted essential gene, however these events are rare so that typically, drive invasiveness is increased substantially (Adolfi et al., 2020; Hou et al., 2024). In addition to the fitness models considered here implying fractional reductions in fertility and pupation rates for single-locus population replacement GD and GDBI, the specific mating fitness may alter the GD dynamics as well. This is particularly the case for a GD which produces heterozygote disadvantage. Either HomeR or ClvR should produce highly stable population modification with homing and resistance rates observed for *Ae. aegypti* GD.

## Conclusions

Our results demonstrate a proof-of-principle of continuous population replacement using low threshold CRISPR/Cas9-based GD in *Ae. aegypti*. Two GD, *nanos*- GD and *zpg*- GD, increased in frequency throughout independent small cage trials throughout 12 generations, with GD carrier frequency reaching 62-68% in populations harboring the *nanos*- GD, and 56-79% in populations with the *zpg*- GD. Gene drive behavior in small cage populations was dependent upon the Cas9 promoter, particularly with respect to GDBI accrual (**Fig. 1**, **Fig. 2**). Both GD caused similar rates of *de novo* mutation introduction at the sgRNA target, but selection led to increased accrual of GDBI in populations harboring the GD with the greater fitness cost. Both population replacement GD produce GDBI at relatively high rates, but are expected to reach and maintain high invasion rates for extended periods of time (**Figure 5**). While performance could be improved using design elements to bias selection in favor of GD, the current *zpg*- GD in particular could be suitable as a robust vehicle for antipathogen effector transgene delivery for sustained arbovirus suppression.

## Methods

### Cage trials

Colonies of *Ae. aegypti* Higg’s White Eye strain and the outcrossed transgenic lines AeaNosC109GD, AeaZpgC109GD, and AeaCFPC109 (here referred to as *nanos*- GD, *zpg*- GD and C109-ECFP, respectively) (Reid et al., 2022) are the sources of the mosquitoes used in this study.

Founder populations of hemizygous males for discrete, non-overlapping cage trials were established separately for each cage trial by outcrossing 20 transgenic males to 100 HWE females for each of the “marker only” (no GD control), *zpg*-GD, and *nanos*-GD lines. This outcrossing step removes those indels which may have accumulated independently during routine maintenance of each GD line. G0 populations of each replicate cage trial were then established at a 1:9 GD male: wild-type (HWE) male release ratio by selecting 15 transgenic male pupae, 135 HWE male pupae and 150 female pupae and introducing separately in 1 cubit-foot cages. Male pupae for both the transgenic line and HWE were reared at the same time and collected on the same day to prevent any mating advantage. Throughout our experiments male pupae were collected earlier than female pupae due to a difference in average developmental time to the pupal stage.

Rearing and counting of individuals for cage trials was performed as described in the following: Mosquitoes larvae were reared in a controlled environment (28° C, 80% humidity, 12 h dark, 12 h light cycle) and fed on tropical fish food flakes (Reid et al., 2022). At pupation, individuals were screened for mCherry eye marker expression. From G1 through G12 of the cage trials, 150 male and 150 female pupae were randomly picked from each respective cohort. The number of individuals showing fluorescent eye marker expression was counted for each sex, and pupae were placed in emergence cups within 1 cubic-foot cages. Adult mosquitoes were supplied with raisins and water *ad libitum*. Eight days following pupation, adult females were fed on defibrinated sheep blood (Colorado Serum Company, CO) using custom-made glass feeders. For the cage trials, females were monitored for blood feeding status and re-fed the next day if fewer than 80% of females had taken a blood meal. Egg papers were placed in cages from days 2-6 post-blood feeding. After removal, eggs were allowed to mature for >4 days in drying conditions prior to hatching under vacuum. For G1 to G12, mosquitoes were reared by hatching eggs from the previous generation, and randomly selecting 500 larvae for rearing at ∼125 individuals per tray (at a density of ∼250 larvae per liter). At second instar, 100 larvae from each replicate were randomly selected and pooled in order to measure GDBI.

### Genotyping assay

Genotyping was performed from G1 through G12 with an allele specific PCR test on randomly sampled male GD carriers from each replicate. Males were screened for the presence of the mCherry eye marker, and ∼10 eye marker positive males per cage were frozen at - 20° C. DNA was extracted from individual carcasses using the Quick DNA Miniprep Plus kit (Zymogen) and eluted into Buffer EB (Qiagen). In the PCR assay 3 oligo-primers were used, which produced differently sized amplicons for the native Carb109 locus (592 bp) and the locus when harboring the GD insertion (336 bp). Results were combined for the 3 replicates of the *nanos*- and *zpg*- GD populations to estimate the allele frequencies over time. PCR conditions were as follows: template genomic DNA, was added to GoTaq green master mix (Promega) with 6.7 picomoles primer F1_gty, 3.3 picomoles primer F2_gty and 10 picomoles primer R_gty. Thermocycling conditions were the following: 95° C for 1 minute, followed by 36 cycles of 95° C for 30 sec, 55° C for 20 sec, 72° C for 40 seconds; with a final extension for 2 minutes at 72° C. PCR products and 100 bp plus ladder (GoldBio) were loaded into 1.7% agarose gels and electrophoresed at 90V for 1 hour.

Gels were stained with ethidium bromide and imaged on a FluorChem Q (Cell Biosciences). The PCR primers for genotyping were:

F1_gty (C109 no transgene/ BR-724 no adaptor): TCGCACCTAATCAGACAGTCG

F2_gty (C109 w/ GD insertion): GAGCAGAGGCAAGAGTAGTG

R_gty (BR-725 no adaptor): CCTGCCTTCATTAAGCTCTTTG

For each generation and GD, the Hardy-Weinberg equilibrium frequency was calculated from the combined observations from triplicate cage trials including those individuals that were homozygous for the recessive allele (i.e. no fluorescent marker allele). The set of equations p+q=1 and 1=q^2^+pq+p^2^ was then solved for the expected frequencies of GD (dominant) and non-GD (recessive) alleles, and the associated genotype frequencies, under Hardy-Weinberg equilibrium conditions. To calculate the averaged allele frequencies in cage trial populations harboring the *nanos*-GD and *zpg*- GD (**Fig. 1D**), a second order polynomial function was fit to the homozygosity assessments based on the 3-primer PCR assays and averaged for 2 generations at a time in order to reduce noise from small sample sizes (**Fig. S3**).

### Quantitation of indels

We measured allele frequencies around the target locus by paired-end sequencing of the target PCR amplicon using total DNA obtained from pools of 100 larvae per cage and generation. A 537 bp amplicon spanning the sgRNA target site of each GD was sequenced from paired-end reads of 300 bp using an Illumina MiSeq instrument, which produced an average of 177,000 paired-end reads per sample. The minimum read coverage for any sample was 13,481 paired-end reads, equivalent to 67 times read coverage of the target amplicon. In total, amplicons from 7200 individual mosquitoes originating from 72 separate pooled samples that were collected from the six GD cage populations were included in the analysis. The 100 second instar larvae (per cage and generation) were selected at random from the ∼500 individual larvae reared during each generation for each cage. As a control 100 larvae pools were collected from the parent HWE line. In order to test for the presence of GDBI in another mosquito population (apart from HWE) capable of transmitting arboviruses, pooled samples of 100 larvae were also harvested from the Orlando strain of *Ae. aegypti* (Dong et al., 2016; Stephenson et al., 2021). The total DNA was extracted from larval pools using Quick DNA Miniprep Plus kit (Zymogen) and eluted into Buffer EB (Qiagen).

The primers utilized to produce the target amplicon bracket the sgRNA target site by >250 bp in both 3’ and 5’ directions. The primers produce a 537 bp amplicon around the C109 locus, when indels are absent. The target amplicons were produced using Q5 High Fidelity PCR polymerase (NEB #M0491) on a Bio-Rad C1000 Touch Thermocycler using a 2-step PCR protocol: 10 cycles with 10 seconds of DNA denaturation at 98° C, 20 seconds primer annealing at 56° C and 30 seconds extension at 72° C; followed by 20 cycles with the annealing temperature increased to 69° C. The PCR products were analyzed on 1.5% agarose gels; bands were confirmed via ethidium bromide staining and the gel portion showing a band signal between 450 bp and 750 bp was excised. The DNA was purified and concentrated from agarose gels using the Nucleospin Gel and PCR Clean-up kit (Takara Bio) and samples were normalized to 1ng/μl using a Qubit fluorometer (Thermo Fisher Scientific). Second round PCR assays using unique dual index sequencing primers and sequencing of resulting PCR amplicons were performed by the University of Missouri Genomics Technology Core. Amplicons were sequenced on the Illumina MiSeq platform, producing 300 bp paired-end reads (PE300).

The resulting paired-end reads were trimmed using Trimmomatic (Bolger et al., 2014). Indel analysis was performed on the trimmed reads using Crispresso2 (Pinello et al., 2016; Clement et al., 2019), with the minimum average read quality set to 10 and GGATAGCCGAAGAAAAGCCA defined as the C109 target specific crRNA sequence. Substitutions were ignored as part of the indel analysis, as multiple single-nucleotide polymorphisms (SNPs) were detected in controls 3’ of the sgRNA target site. GDBI were quantified for each pool as the percentage of reads with insertions or deletions within the sgRNA target site. For the quantitation of *de novo* indels, only mutations which were observed in > 0.5% of all read counts in a sample were included. This value was chosen based on the reasoning that each sample of 100 larvae contains a maximum of 200 non-GD alleles at the C109 locus.

1st round PCR primers for Illumina MiSeq, quantitation of indels (underlined portion is the target amplicon specific primer):

F1_C109amplicon:

ACACTCTTTCCCTACACGACGCTCTTCCGATCT<u>ATGGAAACAAAACACAAGGCATACA</u>

R1_C109amplicon:

GTGACTGGAGTTCAGACGTGTGCTCTTCCGATCT<u>TTATCCGACGAAAATGTGTTCACTG</u>

### Modeling

Modeling studies were performed using the MGDrivE and MGDrivE2 packages with R studio (Sanchez et al., 2020; Wu et al., 2021; R Core Team 2021). The GD cube function is a modified version of the cubeHomingDrive (Cube-CRISPR2MF.R) for a population replacement, homing endonuclease GD with non-negligible resistance allele formation rates (Reid et al., 2022). In this model, individual resistance alleles arising from NHEJ repair of CRISPR/Cas9 catalyzed double-strand DNA breaks at the targeted intergenic locus are assumed to have the same fitness. The GD model includes sex-specific homing rates, resistance allele formation, and maternal deposition rates. The models shown in this paper were parameterized with *nanos*- and *zpg*- GD specific values for homing, resistance allele formation and maternal deposition which were previously determined from extensive empirical data based on single-crosses (Reid et al., 2022).

In the model, six different genotypes resulting from combinations of GD (“H”), wild-type (“W”) and resistance allele or GDBI (“R”) are possible. Fitness cost was implemented in MGDrivE using the cubeModifiers function from the MGDrivE2 package and inputting genotype specific parameters of xiF, xiM, and s; where xiF = female specific pupation rate, xiM= male specific pupation rate, and s= fertility (Wu et al., 2021). The xiF and xiM parameters aggregate reductions in larval and pupal survival rates. In this scenario the pupation rate parameters may also represent an aggregate fitness cost which accounts for the fraction of zygotes of the specified genotypes that contribute genetic material to the next generation in a deterministic population simulation. For the models with a generalized GD fitness cost, the values for the “WH”, “HH” and “HR” genotypes were set to less than 1 in the cubeModifiers function. The default genotype-specific fitness is set to 1 for no fractional reduction in fitness. The fit of the GD models parameterized with different fitness costs was compared, using the observed cage trial allele frequencies as a reference, by calculating the mean absolute scaled error (Hyndman and Koehler, 2006).

## Abbreviations

GD: gene drive
HEG: homing endonuclease gene
HDR: homology-directed repair
NHEJ: non-homologous end-joining
C109: intergenic locus located 3’ of the carboxypeptidase A gene on the 3^rd^ chromosome of Ae. Aegypti
GDBI: gene drive blocking indel mutation
HWE: Higg’s White Eye laboratory strain of Ae. Aegypti
DSB: DNA double-strand break

## Notes

### Competing Interest Statement

The authors have declared no competing interest.

