## Supplemental data for "Performance of low-threshold, population replacement gene drives in cage populations of the yellow fever mosquito, *Aedes aegypti*"

**Tables S1A-C. GD carriers by sex** for the **A.** *nanos*- GD and **B.** *zpg*- GD from combined triplicate cage trials **C**. Fisher's exact test results for sex bias in GD inheritance.

**A.**

| | GD | no GD |
|---|---|---|
| AF35G1 male | 57 | 393 |
| AF35G1 female | 35 | 415 |
| AF35G2 male | 127 | 323 |
| AF35G2 female | 95 | 355 |
| AF35G3 male | 210 | 240 |
| AF35G3 female | 164 | 286 |
| AF35G4 male | 203 | 247 |
| AF35G4 female | 203 | 247 |
| AF35G5 male | 234 | 216 |
| AF35G5 female | 238 | 212 |
| AF35G6 male | 232 | 218 |
| AF35G6 female | 216 | 234 |
| AF35G7 male | 251 | 199 |
| AF35G7 female | 237 | 213 |
| AF35G8 male | 253 | 197 |
| AF35G8 female | 257 | 194 |
| AF35G9 male | 263 | 187 |
| AF35G9 female | 255 | 195 |
| AF35G10 male | 273 | 177 |
| AF35G10 female | 263 | 187 |
| AF35G11 male | 289 | 161 |
| AF35G11 female | 284 | 166 |
| AF35G12 male | 281 | 169 |
| AF35G12 female | 311 | 139 |

**B.**

| | GD | no GD |
|---|---|---|
| AF46G1 male | 101 | 349 |
| AF46G1 female | 57 | 393 |
| AF46G2 male | 108 | 342 |
| AF46G2 female | 113 | 337 |
| AF46G3 male | 168 | 282 |
| AF46G3 female | 154 | 296 |
| AF46G4 male | 195 | 255 |
| AF46G4 female | 187 | 263 |
| AF46G5 male | 187 | 263 |
| AF46G5 female | 187 | 263 |
| AF46G6 male | 204 | 246 |
| AF46G6 female | 199 | 251 |
| AF46G7 male | 198 | 252 |
| AF46G7 female | 202 | 248 |
| AF46G8 male | 187 | 263 |
| AF46G8 female | 211 | 239 |
| AF46G9 male | 225 | 225 |
| AF46G9 female | 233 | 217 |
| AF46G10 male | 276 | 174 |
| AF46G10 female | 267 | 183 |
| AF46G11 male | 270 | 180 |
| AF46G11 female | 284 | 166 |
| AF46G12 male | 276 | 174 |
| AF46G12 female | 298 | 152 |

AF-35 = *Ae. aegypti* carrying *nanos*- GD transgene; AF-46 = *Ae. aegypti* carrying *zpg*- GD transgene.

**C.**

| Generation | **p-value**, *nanos-* | **p-value**, *zpg-* |
|---|---|---|
| G1 | 0.0204 | 0.0002 |
| G2 | 0.0164 | 0.7568 |
| G3 | 0.0023 | 0.366 |
| G4 | >0.9999 | 0.6369 |
| G5 | 0.8413 | >0.9999 |
| G6 | 0.3173 | 0.7886 |
| G7 | 0.3844 | 0.8405 |
| G8 | 0.8403 | 0.1226 |
| G9 | 0.6369 | 0.6407 |
| G10 | 0.5411 | 0.5857 |
| G11 | 0.7816 | 0.3731 |
| G12 | 0.0415 | 0.1452 |

**A.**

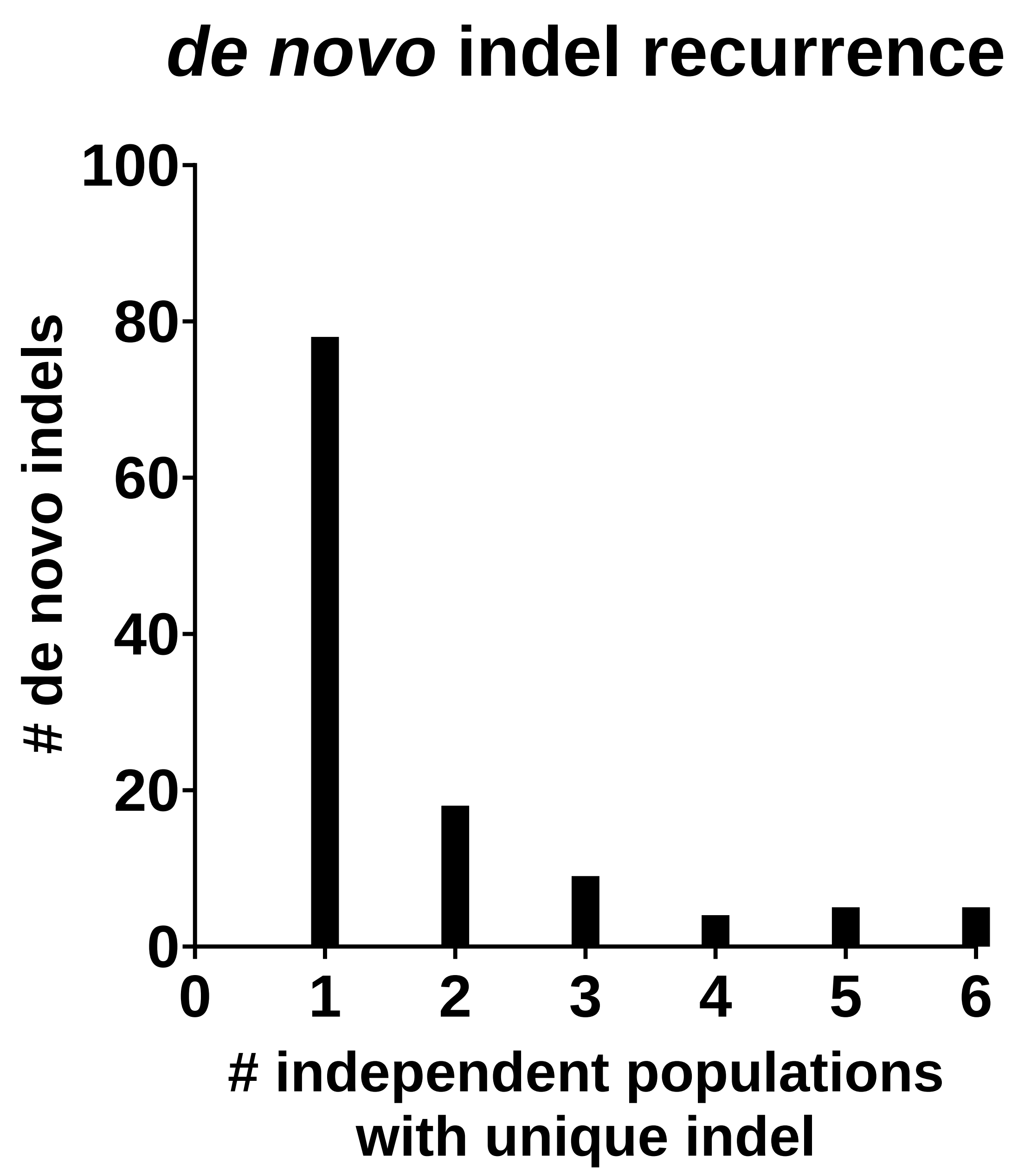


**B.**

| Most frequent indels | Avg. generation first observed | deletion length |
|---|---|---|
| ATCCAACTCCCTGACCCT------TCTTCGGCATATCTAT | 4.0 | 6 bp |
| ATCCAACTCCCTGACCCT----TTTCTTCGGCATATCTAT | 2.7 | 4 bp |
| ATCCAACTCCCTGACCCT---------TCGGCATATCTAT | 5.5 | 9 bp |
| ATCCAACTCCCTGACCCT-----TTCTTCGGCATATCTAT | 7.0 | 5 bp |
| ATCCAACTCCCTGACCC------------GGCATATCTAT | 7.5 | 12 bp |

**Figure S1.** Recurrent *de novo* indels **A)** Observations of recurring indels in non-overlapping cage trial populations. **B)** Five recurring *de novo* mutations were observed in all six independent populations. Three out of five indels, which were identified in all six independent populations occurred within six generations on average. The most common GDBI were short, semi-random deletions (<15 bp) indicative of a predominant NHEJ repair of Cas9 induced double-strand breaks.

**Table S2. *De novo* indels observed in cage trial populations.**

| # | *de novo* indel | population with indel | generation first observed in population |
|---|---|---|---|
| 1 | ATCCAACTCCCTGACCC---------------ATATCTAT | AF35n1,n3,AF46n1 | G2,G9,G4 |
| 2 | ATCCAACTCCCTGAC--------------GGCATATCTAT | AF35n1,n2,n3,AF46n3 | G2,G2,G12,G3 |
| 3 | ATCCAACTCCCTGACCCT----TTTCTTCGGCATATCTAT | AF35n1,n2,n3,AF46n1,n2,n3 | G2,G1,G5,G2,G3,G3 |
| 4 | ATCCAACTCCCTGACCCTGG-----------CATATCTAT | AF35n1,n2,n3,AF46n1,n3 | G2,G1,G2,G4,G5 |
| 5 | ------------------------CTTCGGCATATCTAT | AF35n1 | G2 |
| 6 | ATCCAACTCCCTGACCCT---------TCGGCATATCTAT | AF35n1,n2,n3,AF46n1,n2,n3 | G2,G3,G3,G10,G11,G4 |
| 7 | ATCCAACTCCCTGACC--------TCTTCGGCATATCTAT | AF35n1,n3 | G2,G4 |
| 8 | ATCCAACTCCCTGACC-----TTTTCTTCGGCATATCTAT | AF35n1,n2,n3 AF46n1 | G2,G5,G4,G1 |
| 9 | ATCCAACTCCCTGACCCT------TCTTCGGCATATCTAT | AF35n1,n2,n3,AF46n1,n2,n3 | G4,G3,G4,G7,G3,G3 |
| 10 | ATCCAACTCCCTGACCCTGG--------CGGCATATCTAT | AF35n1,n2,n3 | G5,G11,G12 |
| 11 | ATCCAACTCCCTGACC----------------ATATCTAT | AF35n1,AF46n2 | G5,G8 |
| 12 | ATCCAACTCCCTGACC-------------GGCATATCTAT | AF35n1,n2,n3 | G5,G11,G8 |
| 13 | ATCCAACTCCCTGACCCTGG------GTCAGGG--CTAT | AF35n1 | G5 |
| 14 | ATCCAACTCCCTGACCCTGGGTGTAACTTTTCTTCGGCAT | AF35n1,n2 | G6,G11 |
| 15 | ATCCAACTCCCTGACCCTGGCCTTTTCTTCGGCATATCTA | AF35n1,n3 | G6,G6 |
| 16 | ATCCAACTCCCTGACCC--------CTTCGGCATAACTAT | AF35n1 | G6 |
| 17 | ATCCAACTCCCTGACCCT---------TCGGCATAACTAT | AF35n1,n3,AF46n3 | G6,G2,G7 |
| 18 | ATCCAACTCCCTGACCCT-------CTTCGGCATAACTAT | AF35n1 | G7 |
| 19 | ATCCAACTCCCTGACCC------------GGCATAACTAT | AF35n1 | G7 |
| 20 | ATCCAACTCCCTGACCCTGGCTTT-CTTCGGCATATCTAT | AF35n1,n2,n3,AF46n1,n3 | G7,G5,G9,G5,G5 |
| 21 | ATCCAACTCCCTGACCCTGG-----------CATAACTAT | AF35n1,n2,n3,AF46n2,n3 | G7,G10,G3,G3,G10 |
| 22 | ATCCAACTCCCACACATTGT-------------------- | AF35n1 | G8 |
| 23 | ATCCAACTCCCTGACCCT-----TTCTTCGGCATAACTAT | AF35n1,AF46n3 | G9,G10 |
| 24 | ATCCAACTCCCTGACCCTGG-GCATCTTCGGCATAACTAT | AF35n1 | G9 |
| 25 | ATCCAACTCCCTGACCCTCCCTTT-CTTCGGCATAACTAT | AF35n1 | G9 |
| 26 | AT------------------------------ATAACTAT | AF35n1 | G10 |
| 27 | ATCCAACTCCCTGAC------------------------- | AF35n1 | G10 |
| 28 | ATCCAACTCCCTGACCCT-----TTCTTCGGCATATCTAT | AF35n1,n2,n3,AF46n1,n2,n3 | G11,G3,G4,G11,G9,G4 |
| 29 | ATCCAACTCCCTGACC----------TTCGGCATATCTAT | AF35n1,n2,n3,AF46n1,n3 | G11,G5,G7,G10,G7 |
| 30 | ATCCAACTCCCTGACCCTC----TTCTTCGGCATATCTAT | AF35n1,n3 | G11,G7 |
| 31 | ATCCAACTCCCTGACCC------------GGCATATCTAT | AF35n1,n2,n3,AF46n2,n3 | G11,G7,G5,G3,G9,G10 |
| 32 | ATCCAACTCCCTGACCCTGG-------TCAGGGCATATCT | AF35n1,n3,AF46n2,n3 | G11,G4,G3,G4 |
| 33 | ATCCAACTCCCTGACCCTGGCT---CTTCGGCATAACTAT | AF35n1,n3 | G11,G8 |
| 34 | ATCCAACTCCCTGACCCTGGCATATC---GGCATATCTAT | AF35n1,n3,AF46n1 | G11,G8,G8 |
| 35 | TCCAACTCCCTGACCTCTGACCCTGACCCTCTTCGGCATA | AF35n1,n2 | G11,G7 |
| 36 | ATCCAACTCCCTGACCCT----TTTCTTCGGCATAACTAT | AF35n1 | G12 |
| 37 | ATCCAACTCCCTGAC--------TTCTTCGGCATAACTAT | AF35n1 | G12 |
| 38 | ATCCAACTCCCTGACCCTGGCTCTTTTCTTCGGCATAACT | AF35n1 | G12 |
| 39 | ATCCAACTCCCTGACCCTGAC----------CATATCTAT | AF35n2 | G2 |
| 40 | ATCCAACTCCCTGAC---------TCTTCGGCATATCTAT | AF35n2 | G2 |
| 41 | ATCCAACTCCCTGACCCTGACTTTTCTTCGGCATATCTAT | AF35n2,n3,AF46n3 | G3,G12,G5 |
| 42 | ATCCAACTCCCTGACC----------TTCGGCATAACTAT | AF35n2,n3,AF46n1,n2,3 | G3,G3,G7,G2,G10 |
| 43 | ATCCAACTCCCTGACCCTGG----------GCATATCTAT | AF35n2,n3,AF46n2 | G3,G6,G9 |
| 44 | ATCCAACTCCCTGAC--------TTCTTCGGCATATCTAT | AF35n2,AF46n1 | G5,G3 |
| 45 | ATCCAACTCCCTGACCCT-------CTTCGGCATATCTAT | AF35n2,AF35n3,AF46n1,n2 | G5,G9,G5,G10 |
| 46 | ATCCAACTCCCTGACCCTGGGATCTAT-CGGCATATCTAT | AF35n2 | G5 |
| 47 | ATCCAACTCCCTGATTCTGGCTCTTCTTCGGCGTATCTG- | AF35n2,AF46n2 | G6,G9 |
| 48 | ATCCAACTCCCTGAC-----------TTCGGCATATCTAT | AF35n2 | G7 |
| 49 | ATCCAACTCCCTGACCCTGGCA------------ATCTAT | AF35n2 | G7 |
| 50 | ATCCAACTCCCTGACCCTGG-TTTTCTTCGGCATATCTAT | AF35n2 | G7 |
| 51 | ATCCAACTCCCTGACCCT------TCTTCGGCATAACTAT | AF35n2 | G7 |
| 52 | ATCCAACTCCCTGACCCTGGCTT--CTTCGGCATAACTAT | AF35n2 | G8 |
| 53 | ATCCAACTCCCTGACCCTGG------GTCAG--------- | AF35n2 | G8 |
| 54 | ATCCAACTCCCTGAC-----------------ATATCTAT | AF35n2,AF46n1,n3 | G10,G12,G5 |
| 55 | ATCCAACTCCCTGACCCT--------------------AT | AF35n2 | G12 |
| 56 | ATCCAACTCCC----------------------------- | AF35n3 | G2 |
| 57 | ATCCAACTCCCTGACCC----------------------- | AF35n3 | G2 |
| 58 | ATCCAACTCCCTGACCCT----------CGGCATAACTAT | AF35n3 | G2 |
| 59 | ATCCAACTCCCTGACCCTGG-------------------- | AF35n3,AF46n2 | G3,G12 |
| 60 | ATCCAACTCCCTGACCCTGGCTCGAGATTGCATCGGCATA | AF35n3 | G4 |

| # | *de novo* indel | population with indel | generation first observed |
|---|---|---|---|
| 61 | ATCCAACTCCCTGACCCTGG----TCTTCGGCATATCTAT | AF35n3,AF46n1 | G4,G5 |
| 62 | ATCCAACT--------------TTTCTTCGGCATATCTAT | AF35n3,AF46n2,n3 | G5,G3,G10 |
| 63 | ATCCAACTCCCTGACCCTGGCTGCGAGATTTCTTCGGCAT | AF35n3 | G5 |
| 64 | ATCCAACTCCCTGACC-----TTTTCTTCGGCATAACTAT | AF35n3 | G6 |
| 65 | ATCCAACTCCCTGACCCTGCCTT--CTTCGGCATATCTAT | AF35n3 | G6 |
| 66 | ATCCAACTCCCTGACCCT----------CGGCATATCTAT | AF35n3 | G7 |
| 67 | ATCCAACTCCCTGATTCTGGCTCTTCTTCGGCATATCTAT | AF35n3,AF46n2 | G9,G10 |
| 68 | ATCCAACTCCCTGACCCT---------------------- | AF35n3 | G9 |
| 69 | ATCCAACTCCCTGACCCTGG---TTCTTCGGCATAACTAT | AF35n3 | G9 |
| 70 | ATCCAACTCCCTGATTCTGGCTTTTCTTCGGCATAACTAT | AF35n3 | G10 |
| 71 | ATCCAACTCCCTGACCCTGACCCTTTTCTTCGGCATATCT | AF35n3 | G11 |
| 72 | ATCCAACTCCCTGACCCTGC--------CGGCATATCTAT | AF35n3 | G12 |
| 73 | ATCCAACTCC------------------------------ | AF35n3 | G12 |
| 74 | ATCCAACTCCCTG------------------CATATCTAT | AF46n1 | G2 |
| 75 | ATCCAACTCCCTGACCCTTACTTTTCTTCGGCATATCTAT | AF46n1 | G3 |
| 76 | ATCCAACT-------------------TCGGCATATCTAT | AF46n1 | G5 |
| 77 | ATCCAACTCC-------------------GGCATATCTAT | AF46n1 | G6 |
| 78 | ATCCAACTCCC------------------GGCATATCTAT | AF46n1 | G6 |
| 79 | ---------------------------------------- (63 bp deletion) | AF46n1 | G7 |
| 80 | ATCCAACTCCCTGACCCTGACCCTGACTTCTTCGGCATAT | AF46n1 | G8 |
| 81 | ATCCAACTCCCTGACCCTGG-----CTTCGGCATATCTAT | AF46n1,n3 | G9,G4 |
| 82 | ATCCAACTCCCTGACCCTGG--TTTCTTCGGCATATCTAT | AF46n1,n2 | G9,G8 |
| 83 | ATCCAACTCCCTGACCCTGGCTT--CTTCGGCATATCTAT | AF46n1,n3 | G10,G12 |
| 84 | ATCCAACTA---------------------GGGTATCTAA | AF46n1 | G10 |
| 85 | ATCCAACTCCCTGACCC---CTTTTCTTCGGCATATCTAT | AF46n1 | G11 |
| 86 | ATCCAACTCCCTGACCCTGG-------TCGGCATATCTAT | AF46n1 | G12 |
| 87 | ATCCAACTC-----------------TTCGGCATATCTAT | AF46n2 | G5 |
| 88 | ATCCAACTCCCT-----------TTCTTCGGCATATCTAT | AF46n2 | G5 |
| 89 | ATCCAACTCC-TGACCCT----TTTCTTCGGCATATCTAT | AF46n2 | G5 |
| 90 | ATCCAACTCCCTGACCCTGAC----CTTCGGCATATCTAT | AF46n2,n3 | G6,G4 |
| 91 | ATCCAACTCCCTGACCCCTCCTTTTCTTCGGCATATCTAT | AF46n2 | G10 |
| 92 | ATCCAACTCCCTGACCCTGACCTTTTCTTCGGCATATCTA | AF46n2,n3 | G10,G4 |
| 93 | ATCCAACTCCCTGACCCTGGCAG----------------- | AF46n2 | G10 |
| 94 | ATCCAACTCCCTGACCCC------TCTTCGGCATATCTAT | AF46n2 | G11 |
| 95 | ATCCAACTCCCTGACCCT---------------------T | AF46n2 | G11 |
| 96 | ATCCAACTCCCTGACCCT--CTTTTCTTCGGCATATCTAT | AF46n2 | G12 |
| 97 | ATCCAACTCC-----------TTTTCTTCGGCATATCTAT | AF46n3 | G4 |
| 98 | ATCCAACTCCCTGAGC--------TCTTCGGCATATCTAT | AF46n3 | G4 |
| 99 | ATCCAACTCCCTGG-------------TCGGCATATCTAT | AF46n3 | G4 |
| 100 | ATCCAACTCCCTGACCCA----TATCTACGGCATATCTAT | AF46n3 | G4 |
| 101 | ATCCAACTCCCTGACCCTAAATAATTCCTTTCTTCGGCAT | AF46n3 | G4 |
| 102 | ATCCAACTCCCTGACCCTTC--TTTCTTCGGCATATCTAT | AF46n3 | G4 |
| 103 | ----------------------TTTCTTCGGCATATCTAT | AF46n3 | G5 |
| 104 | ATCCAACTCCCTGACCCTGTATATGTAGATATTTCTTCGG | AF46n3 | G5 |
| 105 | ATCCAACTCCCTGACCCTGG--------CGGCATAACTAT | AF46n3 | G6 |
| 106 | ATCCAAC-------------CTTTTCTTCGGCATATCTAT | AF46n3 | G7 |
| 107 | ATCCAACTCCCTGACCCTG--------------------- | AF46n3 | G7 |
| 108 | ATCCAACTCCCTGAC------TTTTCTTCGGCATATCTAT | AF46n3 | G7 |
| 109 | ATCCAACTCCCTGACCCC----TTTCTTCGGCATATCTAT | AF46n3 | G7 |
| 110 | ATCCAACTCCCT----------TTTCTTCGGCATATCTAT | AF46n3 | G7 |
| 111 | ATCCAACTCCCTGAC--------------GGCATAACTAT | AF46n3 | G8 |
| 112 | ATCCAACTCCCTGACCCTC------CTTCGGCATAACTAT | AF46n3 | G9 |
| 113 | ATCCAACTCCCTGACCCTGAACCTTTTCTTCGGCATAACT | AF46n3 | G9 |
| 114 | ATCCAACTCCCTGACCCTGGC---------GCATATCTAT | AF46n3 | G10 |
| 115 | ATCCAACTCCCTGACCCTGGCTTT-CTTCGGCATAACTAT | AF46n3 | G10 |
| 116 | ATCCAACTCCCTGACCCT---TCTTCTTCGGCATATCTAT | AF46n3 | G10 |
| 117 | ATCCAACTCCCTGACCCT-------CTTCGGCATATCTAT | AF46n3 | G10 |
| 118 | ATCCAACTCCCTGACCCCC---------CGGCATATCTAT | AF46n3 | G10 |
| 119 | ATCCAACTCCCTGACCCTGACCCTACTTTTCTTCGGCATA | AF46n3 | G11 |
| 120 | ATCCAACTCCCTGACCCA------TCTTCGGCATATCTAT | AF46n3 | G12 |

Indels which were identified in all six GD harboring *Ae. aegypti* populations are highlighted in yellow. Only those indels which were observed at a frequency of greater than 0.5% of read counts in a sample are included. The analysis does not include any reads which included any nucleotide calls with a QC score of less than 10 around the sgRNA target site. AF35 = *Ae. aegypti* carrying nanos- GD transgene; AF46= *Ae. aegypti* carrying zpg- GD transgene. n1,n2, n3 = population replicate. G = generation.

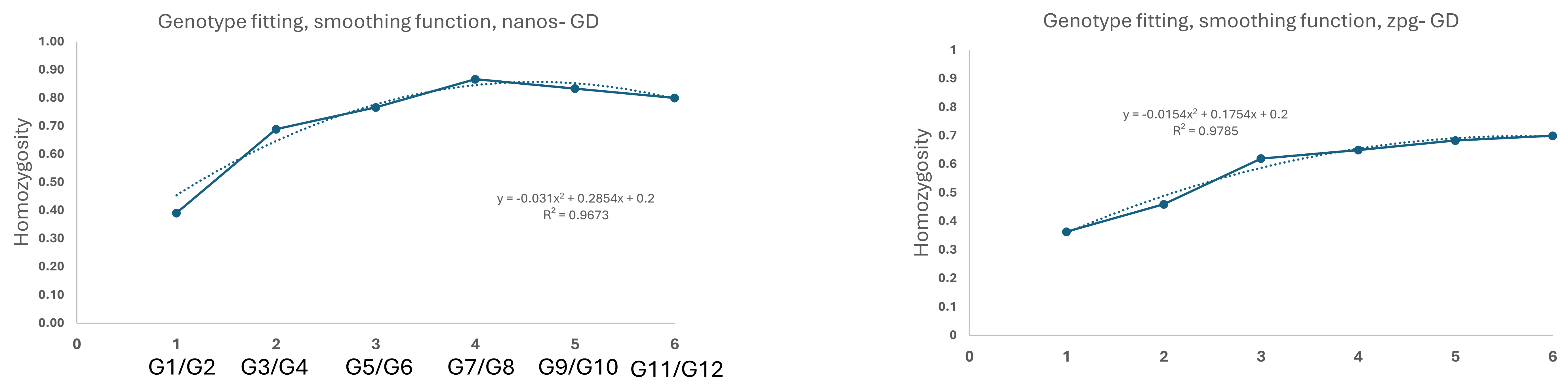


**Figure S3. Genotype smoothing function applied to homozygosity A)** Homozygosity was averaged for two generations at a time for mosquitoes carrying the *nanos*- and *zpg*- GD, respectively, to decrease noise from low sampling numbers (≤30) at each individual generation. A second order polynomial function was fit to the genotype data obtained from allele-specific PCR. The function was applied along with cage trial counts to estimate the GD alleles frequencies shown in Figure 1D.

**Table S3. Fitness parameters and long-term GD inheritance patterns for single-locus, population replacement CRISPR/Cas9 based GD producing *de novo* resistance alleles with inequalities in genotype-specific fitness.**

| **Genotype specific fitness parameters In MGDrivE, xiF=xiM for all runs** | **Fitness Inequality** | **Gene drive inheritance pattern** |
|---|---|---|
| xiM <- c("RR"=.9,"WH"=.9) | **HH=HR=WR>RR=WH** | HH genotype asymptotically approaches fixation |
| xiM <- c("HH"=.9,"RR"=.8,"WH"=.8,"HR"=.9) | **WR>HH=HR>RR=WH** | HH genotype asymptotically approaches fixation |
| xiM <- c("HH"=.9,"RR"=.8,"WH"=.7,"HR"=.8) | **WR>HH>HR=RR>WH** | HH genotype reaches fixation |
| xiM <- c("HH"=.9,"RR"=.8,"WR"=.9,"WH"=.8) | HR>HH=WR>RR=WH | Reaches a stable equilibrium of HH, HR and RR genotypes, with # individuals HR>HH>RR |
| xiM <- c("HH"=.9,"RR"=.8,"WR"=.9,"HR"=.8, "WH"=.9) | **HH=WH=WR>HR=RR** | HH genotype reaches fixation |
| xiM <- c("HH"=.7,"RR"=.6,"WR"=.9,"WH"=.8) | HR>WR>WH>HH>RR | Reaches a stable equilibrium of HH, HR and RR genotypes, with # individuals HR>HH>RR |
| xiM <- c("HH"=.7,"RR"=.6,"WR"=.8,"WH"=.9) | HR>WH>WR>HH>RR | Reaches a stable equilibrium of HH, HR and RR genotypes, with # individuals HR>HH>RR |
| xiM <- c("HH"=.7,"RR"=.6,"WR"=.9,"WH"=.8, "HR"=.7) | **WR>WH>HH=HR>RR** | HH genotype asymptotically approaches fixation |
| xiM <- c("HH"=.7,"RR"=.6,"WR"=.8,"WH"=.9, "HR"=.8) | **WR>WH=HR>HH>RR** | HH genotype asymptotically approaches fixation |

The genotype specific fitness parameters were passed to the cubeModifiers function from MGDrivE2 to model the impact of genotype-specific fitness relationships on GD performance and stability. In addition to the inequalities considered in this table, all combinations with fitness HH AND WH > WR AND RR lead to HH genotype fixation. All inequality combinations with fitness RR AND WR > HH AND WH result in RR genotype fixation. Inequalities with WH=>RR>HH>HR result in fixation of the HH genotype and partial population repression. Inequalities where RR>HH>HR>WH result in RR genotype fixation.

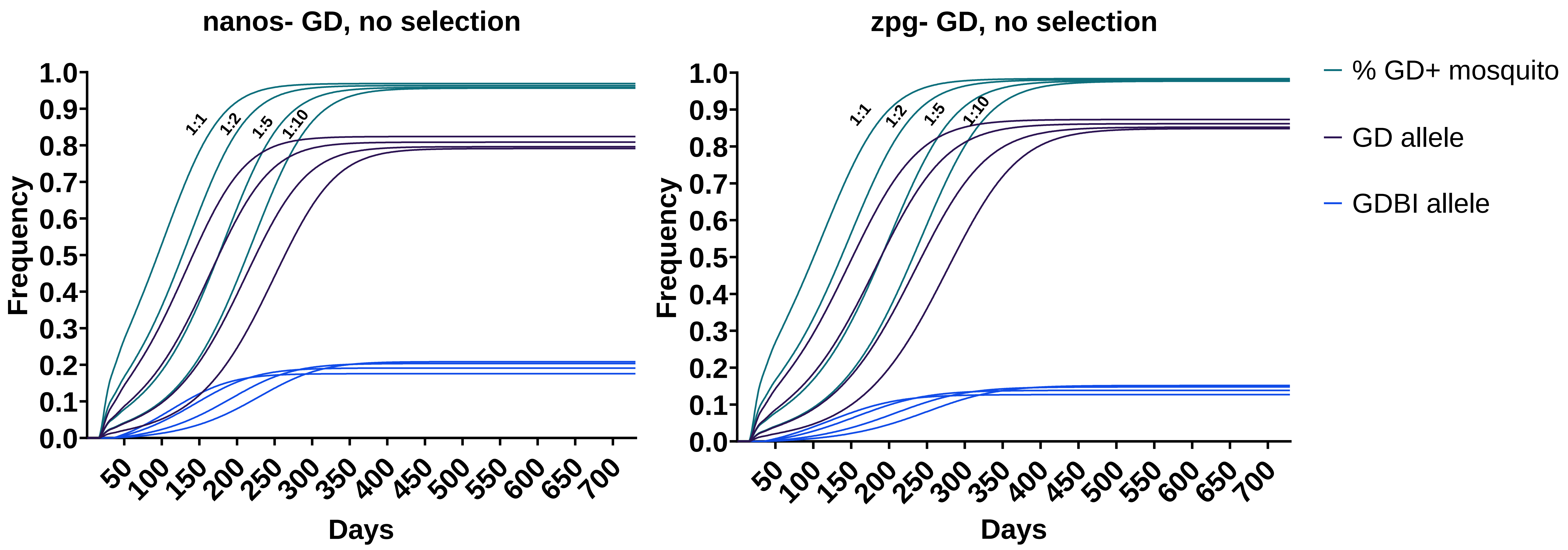


**Figure S4. Predicted behavior of *nanos*- and *zpg*- GD when assuming no fitness costs of GD bearing populations.** Deterministic simulations of low threshold population replacement GD in MGDrivE with varying release conditions of 1:10, 1:5, 1:2 and 1:1 transgenic to wild-type males, using empirically determined parameters for the *Ae. aegypti* nanos- GD and *zpg*- GD. The GD model is designed for a single-locus CRISPR/Cas9 GD with sex-specific gene drive parameters, maternal deposition rates, and gene drive blocking indels (GDBI) having the same fitness as the wild-type. The model is descriptive of a GD targeting an intergenic locus (such as C109), in which the GD transgenes have an associated fitness cost, while GDBI alleles do not incur a fitness cost by default. The release ratio of GD bearing males effects the short term dynamics, with higher release ratios producing maximum GD invasion rates at shorter time scales.

The release ratios do not impact the long term CRISPR/Cas9 GD performance. In the simulations shown, the small differences seen in the equilibrium frequencies result from the release of homozygous GD bearing males into populations with no migration and fixed population size; in models with migration between subpopulations, the equilibrium frequencies would converge.

**A.**

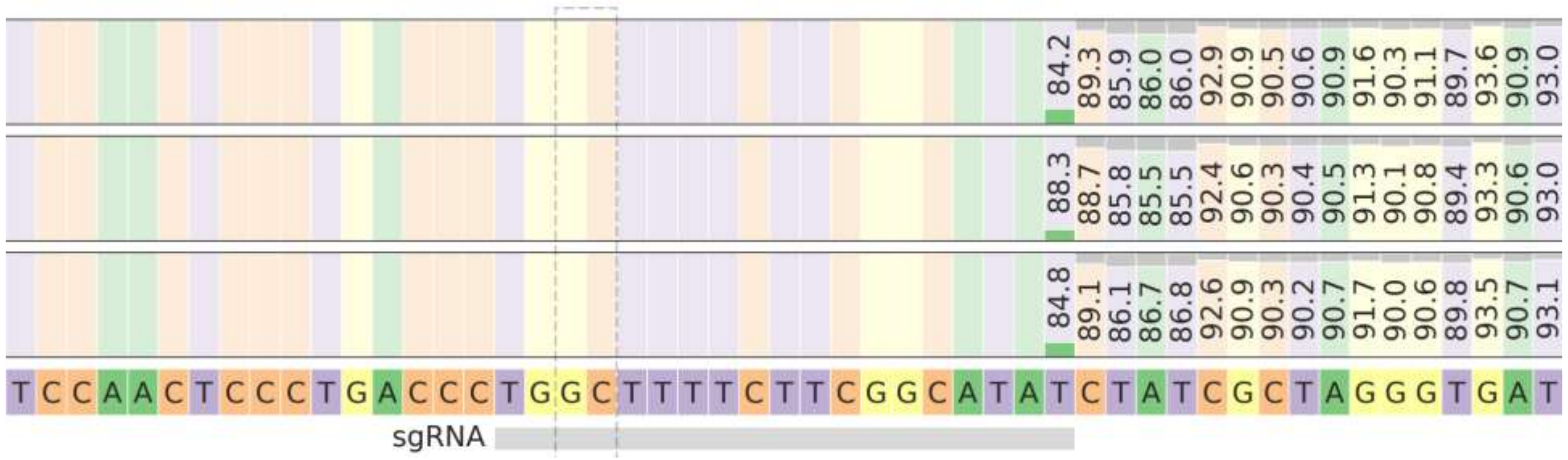


**B.**

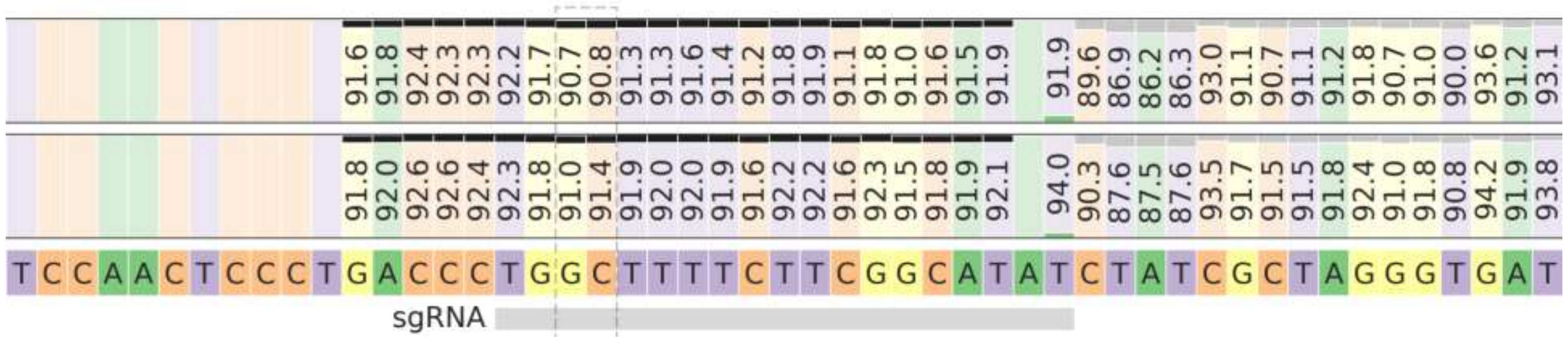


**Figure S5. Target site allele distributions in different *Ae. aegypti* strains A)** Higg's White Eye (HWE) strain and **B)** Orlando strain. The alignment was performed against the Liverpool reference strain, which was originally collected in West Africa in the 1930s, and has a sequence matching the HWE strain proximal at the Cas9 target locus. The HWE strain is derived from the Rexville D (laboratory) strain, which was collected in Bayamon, Puerto Rico in the 1990s. The Orlando strain was established in 1952 from *Ae. aegypti* collections in Orlando, Florida. Around 8-9% of Orlando reads (from replicate samples of 100 individual larvae each) exhibit a single GDBI.
